# Evaluation of phantom doping materials in quantitative susceptibility mapping

**DOI:** 10.1101/2024.02.18.580843

**Authors:** Padriac Hooper, Jin Jin, Kieran O’Brien, Monique Tourell, Simon Daniel Robinson, Markus Barth

**Affiliations:** ARC Training Centre for Innovation in Biomedical Imaging Technology, Brisbane, Australia; Centre for Advanced Imaging, The University of Queensland, Brisbane, Australia; Siemens Healthcare Pty Ltd, Brisbane, Australia; High Field MR Center, Department of Biomedical Imaging and Image-Guided Therapy, Medical University of Vienna, Austria; Christian Doppler Laboratory for MR Imaging Biomarkers, Department for Biomedical Imaging and Image-Guided Therapy, Medical University of Vienna, Austria; School of Information Technology and Electrical Engineering, The University of Queensland, Brisbane, Australia

## Abstract

**Purpose:** To measure magnetic susceptibility (χ) with Quantitative Susceptibility Mapping (QSM) and evaluate its repeatability using four phantom doping materials relevant to QSM applications.

**Methods:** A cylindrical phantom was constructed containing vials of agarose gel doped with two paramagnetic materials (ferritin, USPIO) and two diamagnetic materials (CaCl_2_, CaCO_3_) at five concentrations each. Single orientation QSM measurements (MEDI+0) were carried out on the phantom at 3T and 7T. We measured molar susceptibility (χ_mol_) from QSM and evaluated the test-retest repeatability of χ using the standard error of the measurement (SEM). We evaluated material lifespan by conducting a t-test of χ_mol_ at various timepoints.

**Results:** χ_mol_ (ppm·L·mmol^-1^) were measured as 1.67 ± 0.24 and 0.74 ± 0.09 (USPIO: 3T and 7T, respectively), 10^-2^×(8.13 ± 1.35; 8.13 ± 1.19) (Ferritin: 3T; 7T), 10^-4^×(-2.68 ± 0.24; -2.71 ± 0.37) (CaCl_2_: 3T; 7T), and 10^-5^×(-9.52 ± 1.44; -9.53 ± 1.18) (CaCO_3_: 3T; 7T). The USPIO SEM (1.5 ± 2.0; 5.1 ± 2.0 ppb at 3T; 7T) was greater than the ferritin SEM (1.2 ± 1.0; 2.2 ± 1.3 ppb at 3T; 7T). The CaCl_2_ SEM (7.5 ± 5.5; 1.2 ± 0.6 ppb at 3T; 7T) was greater than the CaCO_3_ SEM (1.2 ± 0.6; 0.9 ± 0.7 ppb at 3T; 7T). We observed no significant changes in molar susceptibility for ferritin and CaCO_3_ over the measured timeframes (24 months and 15 months, respectively).

**Conclusion:** We recommend using ferritin and CaCO_3_ in the construction of susceptibility phantoms, removing later echo times for CaCO_3_ QSM reconstructions.

## 1 Introduction

MRI employs electromagnetic fields to excite and detect nuclear spin resonance. These spins act as probes of the local microenvironment, and with the appropriate model-based analysis, provide a means for quantifying physical properties related to tissue structure and composition. Quantitative parameters (e.g., T_1_, T_2_, diffusion coefficients) can be derived for every voxel in an MRI image and can serve as biomarkers of disease profiles^1,2^. The magnetic susceptibility of tissue, χ, is sensitive to both tissue structure and composition and can be measured using the phase component of the MRI signal acquired from T_2_*-weighted images in Quantitative Susceptibility Mapping (QSM). Clinically, QSM has produced a groundswell of interest, finding applications in mapping calcifications^3^, venous oxygenation^4^, and iron content^5,6^. However, several challenges remain in QSM imaging. Firstly, local χ variations (e.g., air-tissue interfaces) are a source of field distortions and produce regions of low SNR^7^, which lead to artifacts in the susceptibility map^8^. Secondly, the dipole kernel contains zeros in k-space at spatial frequencies corresponding to that of a double cone^9,10^, making dipole inversion an ill-posed problem necessitating regularization. Because of the absence of spatial frequencies at the center of k-space, QSM requires referencing to a known susceptibility value, which is difficult to define in vivo^11^. Additionally, the susceptibility of some tissues are not scalar but tensor, and susceptibility is affected by tissue microstructure, e.g., the radial anisotropy of the myelin sheath^12^.

Susceptibility imaging phantoms that contain uniform regions of known susceptibility are not affected by patient/biological factors^13–16^. As a result, they provide reliable reference values, assess errors in MR acquisition and QSM reconstructions, and enable calibration across scanners and imaging sites^17,18^. In vivo, the predominant cause of tissue susceptibility are iron- or calcium-containing materials that produce paramagnetic (positive) or diamagnetic (negative) susceptibility contrast, motivating the use of iron-based or calcium-based materials, respectively, as χ sources in phantom inclusions. Most phantom inclusions are composed of gel mixtures in place of aqueous mixtures, since gels mimic the in vivo relaxation properties of soft tissues^19^ and embed particles in a fixed position after solidification. Mimicking the signal relaxation and susceptibility of iron- and calcium-based materials provides a realistic evaluation of QSM reconstructions. The protein ferritin, is biologically relevant as a dominant form of iron stored within deep grey matter^20^. To mimic ferritin for QSM applications, Cuna et al. synthesized an iron-filled hydrogel phantom material with variable cluster size, and a comparable molar susceptibility to ferritin *in vivo*^18^. The same research group evaluated QSM susceptibility measurements with a SQUID magnetometer, showing the iron-filled hydrogel phantom material had comparable measurements to those acquired with ultra-high-field scanners^21^. An iron-filled hydrogel phantom material bears a resemblance to iron clustering systems, which is observed Alzheimer’s beta-amyloid plaques^5,6^. Another paramagnetic material of interest is ultrasmall superparamagnetic iron oxide (USPIO), which is applied to QSM as a blood-pool contrast agent^4,22,23^ and in magnetic fluid hyperthermia^24^. USPIO has been used to validate R_2_* mapping at across various field strengths^254^. An important material observed *in vivo* are the insoluble polyhedral salt crystals, calcium carbonate (CaCO_3_), which are present in biochemical analyses within bone mineralization and calcifications^26–28^. Emmerich et al. evaluated CaCO_3_ particles using clinical and ultra-high-field scanners^29^ and used CaCO_3_ particles to study the separation of χ sources in QSM^30^, with *in vivo* comparison to multiple sclerosis lesions^31^. An alternative diamagnetic χ source is calcium chloride (CaCl_2_), which has been used previously by Hopkins et al. to match the susceptibility of bone^32^, and is of practical utility in a phantom due to its high diamagnetism, water solubility and inert chemical properties^33^.

This study aimed to estimate χ and evaluate the repeatability of MR-based χ measurements of two paramagnetic and two diamagnetic materials: USPIO, ferritin (paramagnetic), CaCl_2_ and, CaCO_3_ (diamagnetic). To do this, a cylindrical phantom was constructed containing vials of doped agarose gel. Susceptibility measurements were carried out on the phantom at 3T and 7T, the test-retest repeatability of susceptibility measurements were assessed and evaluated over a 9- to 24-month period. Based on our results we draw conclusions on the suitability of doping materials in QSM phantom studies.

## 2 Methods

### 2.1 Phantom design

Relevant design factors for producing the QSM phantom: it should (i) fit within most RF head coils, (ii) reduce B_0_ and B_1_ inhomogeneities, (iii) provide a means to trap air bubbles, and (iv) be stable over the long-term (≥ 12 months).

The phantom schematics are shown in Figure 1. To ensure fitting in what is probably the most widely used head coil at ultra-high-field currently, the 7 Tesla Nova Medical 1Tx/32Rx head coil (Nova Medical, Wilmington, MA), the outer diameter of the phantom was 150 mm. As most other head coils – including other field strengths – are larger, they would be able to accommodate a phantom of this size. The phantom contains 20 NMR vials (10 mm outer diameter, N-51A, Kimble Glass) of doped agarose gel (details below) surrounded by a solution of ultrapure water (Milli-Q IQ 7000, Merck & Co.), 44.3 weight percentage (wt.%) PVP-40 (PVP-40, Merck & Co.) and 1.7 wt.% NaCl (746398, Merck & Co.) to mimic the electric permittivity and conductivity of white matter at 7T. To reduce B_1_ artifacts and errors during background field correction, vials were positioned at least 10 mm away from the phantom periphery. The phantom was designed to accommodate a bubble trapping compartment outside the Field of View (FOV), connected to the main compartment by a small hole (see Figure 1 a). The tight-fitting non-magnetic closure and an NMR tube cap prevented microbes from entering the phantom and samples, respectively. The modification of electric permittivity in the phantom was important to prevent standing wave artifacts at 7T that would lead to image inhomogeneities.

**Figure 1:**
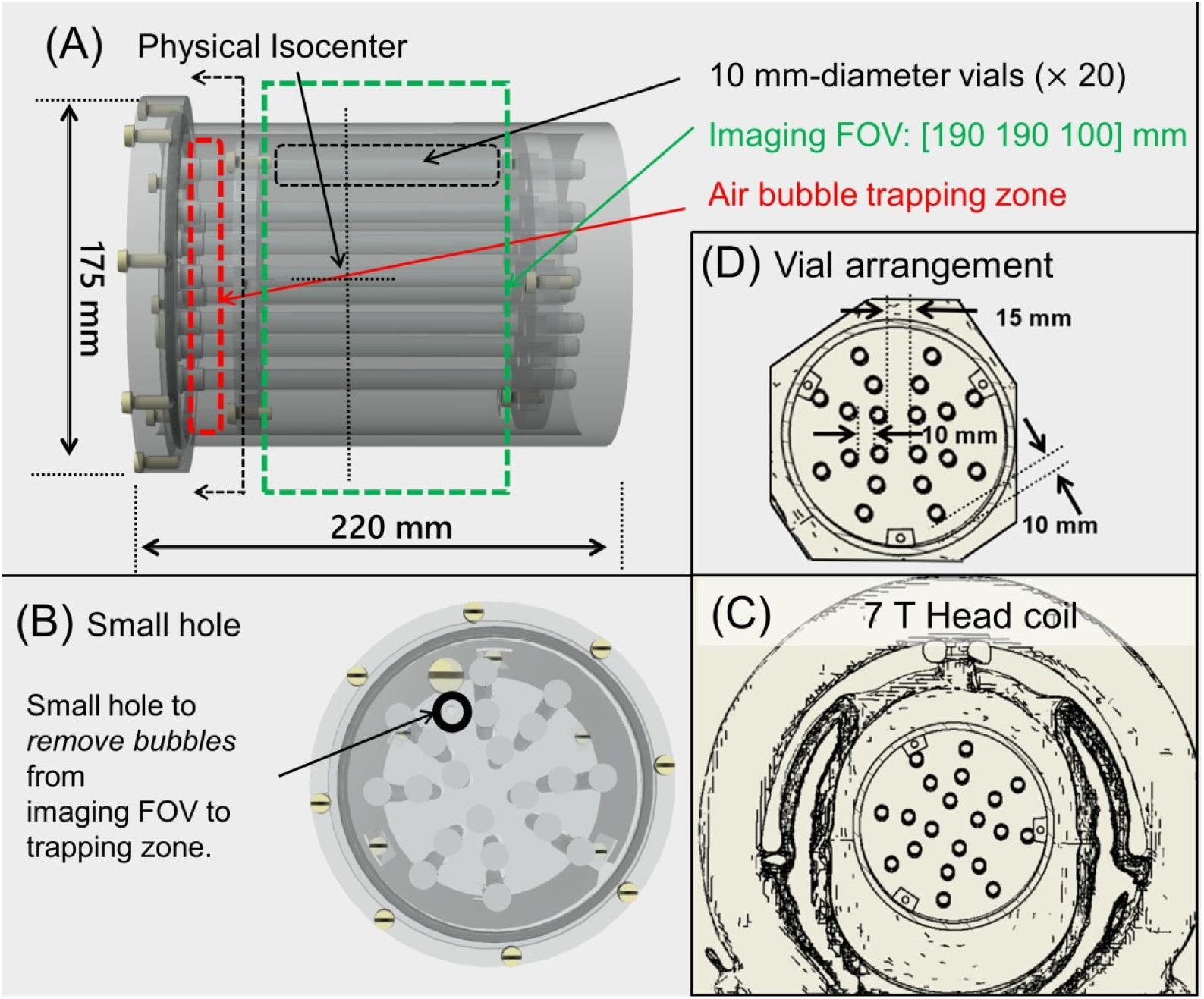
Side (A) and top (B) views of phantom. (C) CAD drawing of the phantom within an approximation of the 7T Nova Medical 32 Rx coil. (D) CAD drawing indicating relative positioning of vials.

The phantom design features four quadrants, each containing five vials. The samples were prepared by doping hot agarose gel with one of the four dopants (USPIO, ferritin, CaCl_2_ and CaCO_3_) at five equally spaced concentrations. Other relevant dopants, Gd-DTPA^34^, hydroxyapatite^35^, and tungsten carbide^30^ were not covered in this study. The concentrations were chosen to more than cover the magnetic susceptibility range expected for in vivo human brain scans, with USPIO ranging from 0.22 to 0.67 mmol/L, ferritin from 3.76 to 10.21 mmol/L, CaCl_2_ from 0.9 to 4.5 mol/L and, CaCO_3_ from 1.0 to 5.0 mol/L. Ultrapure water (Type 1) was used as a solution for the PVP-NaCl mixture to limit the likelihood of microorganisms or magnetic ions entering the samples. The phantom was given at least 48 hours for the PVP-NaCl mixture to settle before scanning^36^.

### 2.2 MR acquisition

The phantom was scanned at two different field strengths of 3T and 7T (all Siemens Healthineers, Erlangen, Germany) with a 20-channel and 32-channel head coil, respectively. Phase and magnitude images were acquired with a multi-echo 3D GRE pulse sequence using the parameters shown in Table 1. We used bipolar acquisitions with shortest possible echo spacings as the results of phase unwrapping are generally more reliable when the shortest possible echo spacings are used^37,38^ because SNR is high and wraps fewer in number^39^. B_0_ shimming was performed using Siemens’ GRE brain sequence and Prescan normalize correction^40^ was used with the 20-channel head coil at 3 T. Images were reconstructed using ASPIRE^41^ for phase and root sum-of-squares for magnitude and converted to NIFTI^42^ format for QSM processing. The phantom was scanned twice at baseline (t_0_), again after 9 months (t_1_), then again twice at 24 months (t_2_). The test-retest scans were acquired within 15 minutes of one another. Some materials were included and removed from the phantom at different timepoints; USPIO and ferritin included at t_0_, t_1_ and t_2_; CaCl_2_ included at t_0_, t_1_; CaCO_3_ included at t_1_, t_2_.

**Table 1:** MR acquisition parameters used in this study.

|  | 3T | 7T |
| --- | --- | --- |
| Model | Magnetom Prisma | Magnetom 7T Plus |
| Gradient readout | Bipolar |  |
| $\text{TE}_1:\Delta\text{TE}:\text{TE}_{\text{max}}$; TR (ms) | 1.87:1.87:22.44; 26 | 3.15:3.15:28.35; 32 |
| Coil | 20-ch Rx head coil | 32-ch Rx head coil |
| Voxel size | 1.0 mm<sup>3</sup> isotropic | 0.7 mm<sup>3</sup> isotropic |
| Acquisition matrix | 192 × 192 × 112 | 272 × 272 × 160 |
| Flip angle; Bandwidth; Averages | 15°; 1000 Hz/pix; 1 | 15°; 340 Hz/pix; 1 |
| Acceleration? type and factor | X | GRAPPA 2 |
| Partial Fourier? factor; Elliptical scanning? | X;X | 6/8 PFA; ✓ |
| Prescan normalize correction? | ✓ | X |

### 2.3 Image pre-processing and corrections

Noise was measured from unfiltered magnitude images as the standard deviation of the intensity in air regions (SD_air_), outside the phantom. Regions of interest (ROIs) 20 × 20 × 160 mm^3^ were drawn manually at the image corners, ensuring ROIs were free of artifacts and matrix borders. SD_air_ has a Rayleigh distribution, which can be corrected to give the expected noise level; SD ^’^= SD/0.66^43^. The signal was measured using cylindrical ROIs (7 mm diameter) drawn manually within each vial, excluding partial volume and/or signal loss at the vial edge, and excluding aliasing artifacts at the 15 distal slices on either end.

The criterion for exclusion data at specific echo time for QSM and R_2_* was set to SNR less than or equal to 10:1. No echo times were excluded for USPIO, ferritin, CaCl_2_; however, for CaCO_3_ echo times longer than 11 ms at 3T and 3 ms at 7T (see Table S1) were excluded. For R_2_* mapping with CaCO_3_ at 7T, echo times longer than 9 ms were excluded. For the statistical analysis of R_2_* mapping, we removed CaCO_3_ concentrations ≥ 4.0 mol·L^-1^ since the SNR was less than or equal to 10:1 at echo times 6 to 9 ms at 7T.

To correct for slight differences in the phantom orientation and positioning between longitudinal acquisitions, the real and imaginary images were manually co-registered (ITK-SNAP v4.0^44^) using the 1^st^ echo magnitude as a reference, interpolating using B-splines. Phase offsets associated with opposite polarity of odd and even echoes due to a bipolar acquisition^45^ were corrected using MCPC-3D-S^37^. χ-induced geometric distortions occur in the readout direction of bipolar GRE acquisitions due to the opposing direction of odd and even readout gradients^46^. Note that geometric distortions were not observed at 3T due to the high readout pixel bandwidth (see Table 1). Field maps of odd and even echoes were created, combining the phase using nonlinear complex fitting^47^, then unwrapped using SPURS^48^. The voxel displacement map was calculated by dividing the field map by the readout pixel bandwidth^49^. The warped field was generated by applying smoothing to the voxel displacement map^50^. The real and imaginary images were unwarped (SPM12)^51^ with tri-linear interpolation. Gibbs artifacts associated with sharp transitions in signal intensity were observed at the vial boundary. A Gibbs ringing correction was applied to the real and imaginary images using sub-voxel shifts in all three spatial dimensions^52,53^.

Zero-padding of the complex GRE data matrix by a factor of 1.5 was used prior to non-linear field map estimation as it is known to result in a more accurate field map^54^. Moreover, it is well established that zero-padding reduces aliasing associated with the Fourier Transform, which affects both background field correction and dipole inversion stages of QSM processing.

### 2.4 R_2_* mapping

R_2_* maps were generated by voxel-wise nonlinear fitting of the multi-echo magnitude image using the Levenberg-Marquardt algorithm. Clump/agglomerate masks were produced as follows: (1) determining local R_2_* outliers (median + 3*IQR within each vial), then (2) finding continuous (6-connectivity) regions of R_2_* outliers larger than 1 mm^3^. From the clump/agglomerate masks, we report (i) the percentage of local R_2_* outliers per vial, and (ii) the range of clump/agglomerate sizes in mm^3^.

### 2.5 QSM

The phase across echo times was fitted using nonlinear complex fitting^47^, then unwrapped using SPURS^48^. Background fields were then corrected using V-SHARP^55^, SMV-radii = 1:1:10 mm (rounding to the nearest integer towards infinity). An ‘initial mask’ was generated by thresholding values within the 1^st^-echo magnitude image greater than 2 % of its maximum intensity. A morphological opening operation, which removes disconnected voxels from the mask, was applied using a structural spherical element of 5 mm radius. The maximum SMV radius was set to 10 mm since the vials positioned close to the perimeter were 10 mm from the mask edge, which is a limitation of V-SHARP^56^. The mask imposed on the field map and on χ during dipole inversion was set as the entire matrix as including the spatial distribution of all frequencies and χ sources improves background field correction^57^. For 7T data, residual RF transmit coil fields (B_1_ ^+^) were corrected by fitting and subtracting a fourth-order 3D polynomial^58,59^.

A ‘reference mask’ (M_reference_) was derived analogous to the so-called ‘CSF mask’ for zero-referencing in MEDI+0 ^60,61^ in 3 steps: (1) R_2_* thresholding at 5 s^−1^, (2) morphological closing (3 mm radius), then (3) morphological erosion (10 mm radius), see Figure S1. The ‘weighting map’ (used to weigh the data consistency term) was computed as described in SEPIA documentation (https://sepia-documentation.readthedocs.io/en/latest/method/weightings.html)^62^. The field noise map was inverted, normalized using the median and upper IQR, re-centered to 1, then global outliers (defined as median + 3*IQR) were replaced with a 3 × 3 × 3 voxel box filtered copy. The relative residual (Equation 3) of S_measured_ (Equation 2) and S_simulated_ (Equation 1) was computed as follows,

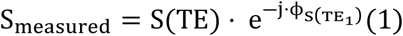

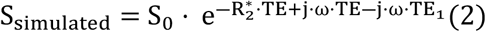

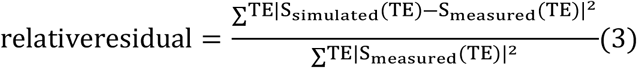

In these equations, ω was the angular frequency determined during field mapping, S_0_ was the extrapolated signal magnitude at TE=0, S_measured_ was the measured data with the phase subtracted from the 1^st^ echo, and S_simulated_ was the simulated mono-exponential model signal with the phase subtracted from the 1^st^ echo. The relative residual map was brought into a weighting component using a threshold of 0.3, which was then used to modulate the weighting map^62^.

For dipole inversion, we used MEDI+0^60,61^ using the default regularization parameters (λ_1_=1000, λ_2_=100). The MERIT parameter was set to false. MEDI with SMV-filtering was not used since it may emphasize the high frequency components more than low frequency components within a susceptibility distribution^8^. The percentage threshold parameter (c_∇_) was optimized by minimizing the streaking artifact with MEDI+0 turned ‘off’, which was quantified as the standard deviation within the reference mask (SD_Ref_)^60,63^. The optimal c_∇_ was found to be 0.5 for both 3T and 7T scans (see Figure S2) and was used for subsequent QSM reconstructions.

The phantom design and vial positioning lends itself to more direct quantification of susceptibility without performing dipole inversion. We used the following analytical model^7^ (referred to as the ‘infinite cylinder model’) to generate a simulated susceptibility map,

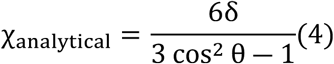

In Equation (4), δ=Δf_local_/f_0_ is the local frequency in Hz (Δf_local_) normalized to the Larmor frequency (f_0_) in parts-per-million (ppm). The angle relative to the applied field vector (θ) is assumed to equal 0°, therefore making χ_analytical_ = 3δ.

### 2.6 Automatic segmentation

We performed automatic segmentation instead of manual segmentation to obtain clear and consistent cylindrical ROIs for statistical analysis. The ‘initial mask’ (refer to section 2.5) was eroded by 10 mm, which is the distance between the distal vials and the mask edge. The complement of the non-eroded reference mask M_reference_ (refer to Figure S1) was multiplied by the eroded initial mask, obtaining a mask of the 10 mm vials. To exclude erroneous voxels at the vial edge, the obtained mask was eroded by 3 mm. We also excluded the 15 distal slices on either end which were prone to aliasing artifacts. Segmentation was performed on the obtained mask using the cluster function within FSL^64^.

### 2.7 Statistical analysis

The molar concentration (c_mol_) was fitted against the mean ROI measurement for χ and R_2_*, respectively, with a least-squares regression to determine the linear fit:

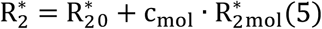

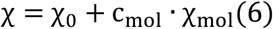

We used robust regression with a bisquare weighting function, which reduces the weight of independent variables with a high least-squares residual. R_2_*_0_; χ_0_ are equal to the agarose R_2_* value; χ value, respectively. R_2_*_mol_; χ_mol_ are equal to the R_2_* relaxivity; molar susceptibility, respectively.

To assess test-retest repeatability of susceptibility measurements, a single-score coefficient of reliability (ICC) was calculated using a two-way ANOVA model with absolute agreement (‘A-1’)^66,67^. We also computed standard error of the measurement (SEM), which is given in terms of the standard deviation of test-retest measurements (SD) and the ICC^17^,

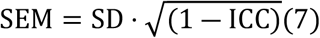

Smaller values of SEM represent greater test-retest precision. To determine material lifespan, we performed a t-test to detect a significant difference between baseline χ_mol_ values to timepoint χ_mol_ value; the first timepoint at which there was a significant change was determined to be the material lifespan. We also performed a t-test to detect a possible correlation between time (t=0, 9, 24 month) and timepoint χ_mol_ values.

Bland–Altman (BA) analysis for repeated measurements per subject (phantom) was applied to evaluate agreement between 3T and 7T scanners^68^. We performed the following BA analyses: (1) to determine the agreement in χ with field strength; ferritin, CaCl_2_ and CaCO_3_), (2) to determine the agreement in the product of the magnetization (M=χ·B_0_) with field strength; USPIO^4^, (3) to determine the agreement in R_2_*/B_0_ with field strength; CaCO_3_ ^31^, (4) to determine the agreement in R_2_* with field strength; USPIO, ferritin and CaCl_2_. We also performed a t-test to detect linear correlation between cross-field measurements; if detected, then we performed linear regression to quantify the trend between cross-field measurements. To validate the dipole inversion step, Bland–Altman (BA) analysis for repeated measurements per subject (phantom)^66^ was applied to evaluate agreement between the simulated analytical susceptibility map (χ_analytical_ from Equation (4)) and the measured susceptibility map (χ_MEDI+0_).

## 3 Results

Maps and plots of susceptibility for the four used materials and five concentrations are shown in Figures 2 to 5. On the scatter plots, the fitted regression was dotted, and, where given, the 95% confidence intervals were solid. The vertical error bars were the standard deviation of the ROI measurement. The Bland-Altman mean difference (bias) line was solid, and the limits-of-agreement were dotted. On the correlation plots, the identity line (y = x) was dotted; the fitted regression and 95% confidence interval lines were solid.

**Figure 2:**
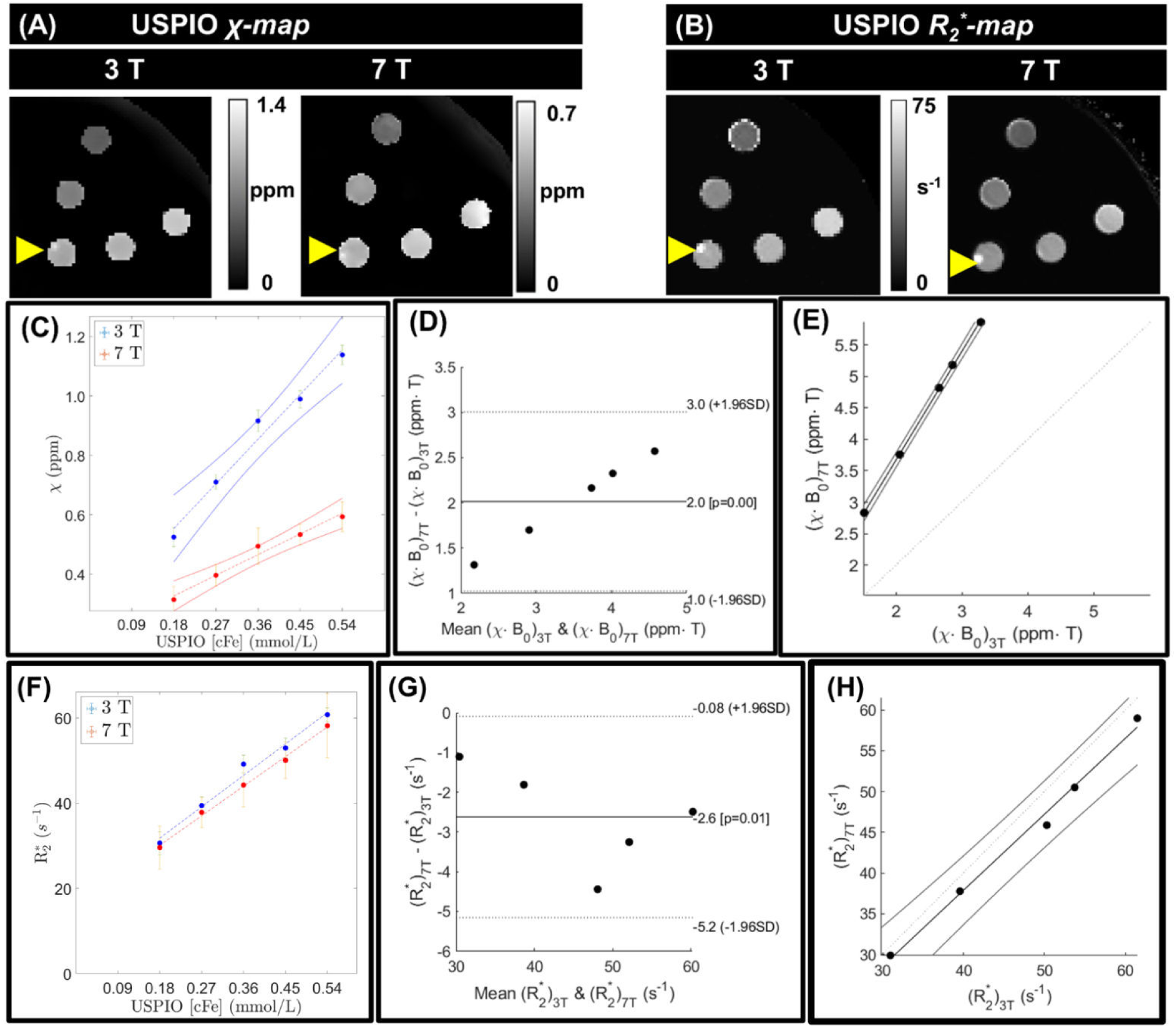
USPIO χ maps (A) and R_2_* maps (B). The windowing of the χ map at 7T was set to half that of 3T. The yellow arrowhead indicated the presence of an agglomerate at USPIO 0.36 mmol/L, manifesting as bright on both the χ and R_2_* maps. (C) Scatter plot of χ as a function of USPIO concentration. (D) Bland-Altman and (E) correlation plots of (χ·B_0_)_3T_ versus (χ·B_0_)_7T_. (F) Scatter plot of R_2_* as a function of USPIO concentration. (G) Bland-Altman and (H) correlation plots of (R_2_*)_3T_ versus (R_2_*)_7T_.

### 3.1 Analysis of clumps/agglomerates and other confounds

We termed “agglomerates” as bright clusters on a susceptibility map and “clumps” as dark clusters on a susceptibility map; clumps/agglomerates each were bright on the R_2_* maps. At 3T, the mean clump/agglomerate volumes of ferritin (3.49 mm^3^) were roughly 6% larger than those of USPIO (3.27 mm^3^). At 7T, the mean clump/agglomerate volumes for ferritin (13.01 mm^3^) were roughly twice those of USPIO (6.40 mm^3^). A single agglomerate was observed in USPIO 0.45 mmol/L (see Figure 2), and several clumps were observed in ferritin 10.2 mmol/L (see Figure 3).

**Figure 3:**
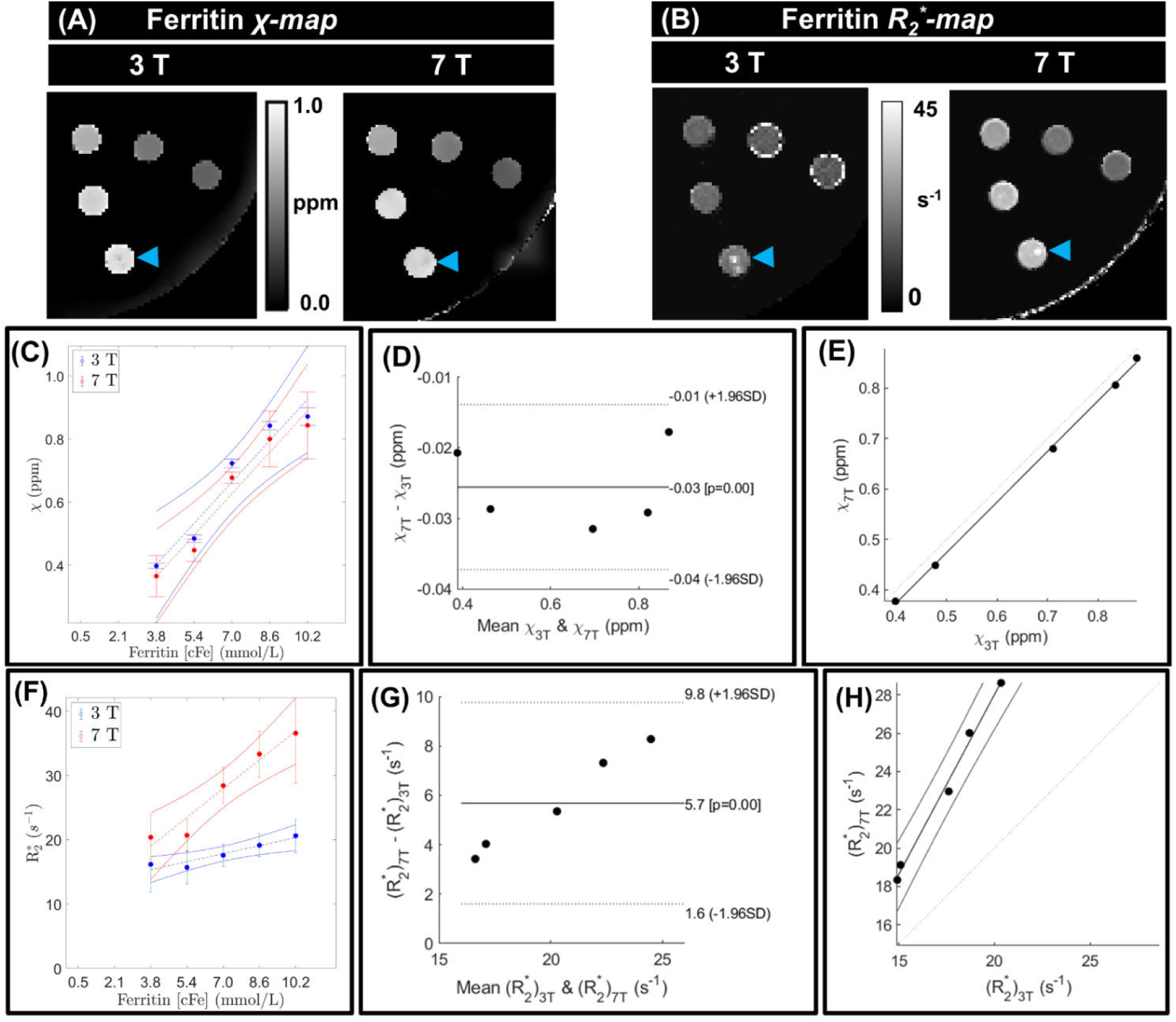
Ferritin χ maps (A) and R_2_* maps (B). At 10.2 mmol/L, ferritin clumps were marked with a blue arrowhead, manifesting as dark on χ (diamagnetic) and bright on R_2_*. (C) Scatter plot of χ as a function of ferritin concentration. (D) Bland-Altman and (E) correlation plots of χ_3T_ versus χ_7T_. (F) Scatter plot of R_2_* as a function of ferritin concentration. (G) Bland-Altman and (H) plots of (R_2_*)_3T_ versus (R_2_*)_7T_.

For CaCl_2_ 4.5 mol/L, we observed heterogeneity, which was attributed to it reaching the water solubility limit^33^; thus, we excluded CaCl_2_ 4.5 mol/L from all statistical analysis. CaCO_3_ concentrations ≥ 4.0 mol/L were heterogeneous on the R_2_* map (see Figure 5) due to low SNR; thus, these two concentrations were excluded from the R_2_* statistical analysis. At 3T, the mean clump/agglomerate volumes for CaCl_2_ (1.46 mm^3^) were roughly 15% larger than CaCO_3_ (1.27 mm^3^). At 7T, the mean clump/agglomerate volumes for CaCl_2_ (5.59 mm^3^) were roughly half that of CaCO_3_ (10.40 mm^3^). The mean number of R_2_* map outlier voxels per vial was less than 3.0 % for each material group at each field strength (see Table S2).

### 3.2 Linear regression (c_mol_ versus χ)

The linearity was excellent, R^2^ > 0.9, for all 4 materials and field strengths. For USPIO, χ_mol_ at 3T (1.67 ppm·mmol^−1^·L) was roughly 2.3 times greater than its χ_mol_ at 7T (0.74 ppm·mmol^−1^·L). For ferritin, χ_mol_ at 3T and 7T were equal (8.13×10^-2^ ppm·mmol^−1^·L). For CaCl_2_, χ_mol_ at 3T (2.68 ×10^-4^ ppm·mmol^−1^·L) was roughly 1% less than its χ_mol_ at 7T (2.71 ×10^-4^ ppm·mmol^−1^·L). For CaCO_3_, χ_mol_ at 3T (9. 52 ×10^-5^ ppm·mmol^−1^·L) was also roughly 1% less than its χ_mol_ at 7T (9.53 ×10^-5^ ppm·mmol^−1^·L).

**Table 2:** Linear fitting coefficients (c_mol_ versus χ) and linearity (R^2^) at 3T and 7T, respectively, from MEDI+0 reconstructed susceptibility maps. Coefficients are reported with standard error.

| | | USPIO | Ferritin | $\text{CaCl}_2$ | $\text{CaCO}_3$ |
| --- | --- | --- | --- | --- | --- |
| $\chi_{\text{mol}}$ ($\text{ppm} \cdot \text{L} \cdot \text{mmol}^{-1}$) | 3T | $1.67 \pm 0.24$ | $(8.13 \pm 1.35) \times 10^{-2}$ | $(-2.68 \pm 0.24) \times 10^{-4}$ | $(-9.52 \pm 1.44) \times 10^{-5}$ |
| | 7T | $0.74 \pm 0.09$ | $(8.13 \pm 1.19) \times 10^{-2}$ | $(-2.71 \pm 0.37) \times 10^{-4}$ | $(-9.53 \pm 1.18) \times 10^{-5}$ |
| $\chi_0$ (ppb) | 3T | $253 \pm 61$ | $80 \pm 81$ | $25 \pm 80$ | $190 \pm 43$ |
| | 7T | $188 \pm 28$ | $53 \pm 91$ | $19 \pm 91$ | $165 \pm 39$ |
| $R^2$ ($c_{\text{mol}}$ versus $\chi$) | 3T | 0.95 | 0.95 | 0.98 | 0.94 |
|  | 7T | 0.95 | 0.94 | 0.97 | 0.96 |

### 3.3 Linear regression (c_mol_ versus R_2_*)

The linearity was poor (R^2^<0.5) for CaCl_2_ at 3T, good (0.75<R^2^<0.9) for CaCl_2_ at 7T and CaCO_3_ at 7T, and excellent (R^2^ > 0.9) for CaCO_3_ at 3T, USPIO and ferritin at both field strengths (see Table S3). For USPIO, R_2_*_mol_ at 7T (77.5 s^-1^·mmol^-1^·L) was roughly 6% less than R_2_*_mol_ at 3T (82.3 s^-1^·mmol^-1^·L). For ferritin, R_2_*_mol_ at 7T (0.77 s^-1^·mmol^-1^·L) was roughly 3.6 times less than R_2_*_mol_ at 7T (2.78 s^-1^·mmol^-1^·L). The linearity of CaCl_2_ was poor at 3T, we report R_2_*_mol_ mean ± standard deviation as 13.1 ± 0.6 s^-1^; 14.0 ± 1.1 s^-1^ at 3T; 7T, respectively. For CaCO_3_, R_2_*_mol_ at 7T (0.869 s^-1^·mmol^-1^·L) was roughly 2.5 times greater than R_2_*_mol_ at 7T (0.352 s^-1^·mmol^-1^·L).

### 3.4 Test-retest repeatability

Coefficient of reliability (ICC) and standard error of the measurement (SEM) of test-retest measurements are shown in table 3. The test-retest repeatability was ‘excellent’ (ICC > 0.90) for each material at each field strength. For ferritin and CaCO_3_, the SEM each were within 5 ppb at both field strengths. At 3T, the USPIO SEM (1.5 ± 2.0 ppb) was 25% larger than the ferritin SEM (1.2 ± 1.0 ppb). At 7T, the USPIO SEM (5.1 ± 2.0 ppb) was 1.8 times larger than the ferritin SEM (2.2 ± 1.3 ppb). At 3T, the CaCl_2_ SEM (7.5 ± 5.5 ppb) was 625 % larger than the CaCO_3_ SEM (1.2 ± 0.6 ppb). At 7T, the CaCl_2_ SEM (1.2 ± 0.6 ppb) was 33% larger than the CaCO_3_ SEM (0.9 ± 0.7 ppb).

**Table 3:** Test-retest repeatability and lifespan of χ measurements for concentrations 1 to 5 of each material group. Besides material lifespan, values given at 3T; 7T, respectively. Coefficient of reliability (ICC) reported as: ICC lower interval to ICC upper interval. Standard error of the measurement (SEM) of test-retest measurements reported as mean ± standard deviation. Material lifespan was based on the t-tests comparing baseline χ_mol_ values to χ_mol_ value at various timepoints.

| | USPIO | Ferritin | $\text{CaCl}_2$ | $\text{CaCO}_3$ |
| --- | --- | --- | --- | --- |
| ICC, test-retest | 0.99-1.00;<br>0.99-1.00 | 0.99-1.00;<br>1.00-1.00 | 0.99-1.00;<br>1.00-1.00 | 1.00-1.00;<br>1.00-1.00 |
| SEM, test-retest (ppb) | $1.5 \pm 2.0$;<br>$5.1 \pm 2.0$ | $1.2 \pm 1.0$;<br>$2.2 \pm 1.3$ | $7.5 \pm 5.5$;<br>$1.2 \pm 0.6$ | $0.8 \pm 0.6$;<br>$0.9 \pm 0.7$ |
| Material lifespan | $\leq 24$ months<br>($p=0.05$ at 3T) | $> 24$ months<br>(no significant change observed) | $\leq 9$ months<br>($p=0.04$ at 7T) | $> 15$ months<br>(no significant change observed) |

### 3.5 Material lifespan

Comparing the molar susceptibility measurements from t_0_ to t_1_ showed no significant change for USPIO (p=0.32; p=0.06 at 3T; 7T), ferritin (p=0.55; p=0.11 at 3T; 7T), CaCl_2_ at 3T (p=0.71), but there was a significant change for CaCl_2_ at 7T (p=0.04), see table 3. Comparing the molar susceptibility measurements from t_0_ to t_2_ showed a significant change for USPIO at 3T (p=0.05) but not at 7T (p=0.10), and no significant change for ferritin (p=0.24; p=0.32 at 3T; 7T). Comparing the molar susceptibility measurements from t_1_ to t_2_ showed no significant change for CaCO_3_ (p=0.38; p=0.12 at 3T; 7T). There was no significant linear correlation between time (t_month_=[0,9,24]) and molar susceptibility (χ_mol_) for USPIO (p=0.38; p=0.82 at 3T; 7T) and ferritin (p=0.28; p=0.70 at 3T; 7T).

### 3.6 Cross-field strength agreement

For both ferritin and CaCO_3_, the χ_3T_ versus χ_7T_ relationship showed a significant bias; -0.03 ppm (p=0.00) for ferritin and -0.02 ppm (p=0.01) for CaCO_3_. For both ferritin and CaCO_3_, the correlation between χ_3T_ versus χ_7T_ was significant (p=0.02), and the correlation line was y=1.00x-0.03 for ferritin, and y=0.99x-0.02 for CaCO_3_. For CaCl_2_, the χ_3T_ versus χ_7T_ relationship showed no significant bias (p=0.50), and no significant correlation (p=0.08). The product of USPIO susceptibility and field strength, χ·B_0_, showed a significant bias of 2.0 ppm·T (p=0.00). The correlation between (χ·B_0_)_3T_ and (χ·B_0_)_7T_ was significant (p=0.02), and the correlation line was y=1.69x+0.29.

The USPIO (R_2_*)_3T_ and (R_2_*)_7T_ showed significant bias of -2.6 s^-1^ (p=0.01). The correlation between (R_2_*)_3T_ and (R_2_*)_7T_ was significant (p=0.02), and the correlation line was y=0.93x+0.55. The ferritin (R_2_*)_3T_ and (R_2_*)_7T_ showed significant bias 5.7 s^-1^ (p=0.00). The correlation between (R_2_*)_3T_ and (R_2_*)_7T_ was significant (p=0.02), and the correlation line was y=1.87x-9.48. The CaCl_2_ (R_2_*)_3T_ and (R_2_*)_7T_ showed no significant bias (p=0.32), and no significant correlation (p=1.00). The quotient of CaCO_3_ R_2_* and field strength, R_2_*/B_0_, showed a significant bias of -1.4 s^-1^·T^-1^ (p=0.01). The correlation between (R_2_*/B_0_)_3T_ and (R_2_*/B_0_)_7T_ was not significant (p=0.33).

### 3.7 Comparison to analytical model

χ_analytical_ and χ_MEDI+0_ showed no significant bias for each material (USPIO, ferritin, CaCl_2_, CaCO_3_) and field strength (3T, 7T) as indicated in Table S4 (p>0.05). The correlation between χ_analytical_ and χ_MEDI+0_ was significant for each material (USPIO, ferritin, CaCl_2_, CaCO_3_) and field strength (3T, 7T) as indicated in Table S4 (p=0.02). At 3T, the correlation lines were y=1.00x+0.00 (USPIO), y=0.99x+0.01 (ferritin), y=0.99x+0.00 (CaCl_2_ and CaCO_3_). At 3T, the correlation lines were y=0.97x+0.01 (USPIO), y=0.98x+0.01 (ferritin), y=0.97x-0.02 (CaCl_2_) and y=0.98x+0.02 (CaCO_3_).

## 4 Discussion

### 4.1 Acquisition and processing

The field-to-susceptibility dipole inversion is formulated as a L_1_-/L_2_-norm regularization; requiring careful tuning of the regularization terms, trading off excessive noise propagation (streaking artifacts) versus the suppression of image features within the noise level^69^. We used MEDI-based QSM reconstruction as the information from magnitude images helped to accurately regularize susceptibility gradients reduceing error propagation during dipole inversion. Moreover, the MEDI algorithm uses a non-linear data fidelity term, which better handles dipole incompatible fields than the linear data fidelity variant^8^.

We observed minor deviations in SNR and local field surrounding the clumps/agglomerates. The mean number of R_2_* map outlier voxels per vial was small (less than 3.0 % for each material group, as indicated in Table S2). The SNR within phase images are higher than SNR within magnitude images^70,71^; which implies that SNR within susceptibility maps are higher than SNR within R_2_* maps. This could explain why the minor deviations in SNR surrounding clumps/agglomerates led to larger deviations in the R_2_* maps than in the susceptibility maps. Errors associated with clumps/agglomerates were mitigated by modulating the weighting map using information from a simulated monoexponential signal (as described in section 2.5).

### 4.2 Test-retest repeatability and material lifespan

A key aim within the quantitative MRI community is the development of phantoms that can assist system stability^72,73^. This requires materials that are stable within minutes of repeated measurements (test-retest measurements), as well as within months (longitudinal measurements). At both field strengths, our test-retest measurements indicated that the ferritin SEM was less than the USPIO SEM; and the CaCO_3_ SEM was less than the CaCl_2_ SEM. Dielectric artifacts are an intrinsic limitation that should be considered when using CaCl_2_ in phantoms. The dielectric artifacts lead to the elevated CaCl_2_SEM in the test-retest measurement. At both 3T and 7T, shading about the CaCl_2_ vials was observed (see Figure 4B), and was attributed to the interaction between CaCl_2_ conductivity and the B_1_ field, which leads to signal attenuations that are unevenly distributed throughout the imaging FOV^74^. For USPIO, the high R_2_* dephasing and the high local fields contributed to the elevated USPIO SEM. For ferritin and CaCO_3_, the SEM each were within 5 ppb at both field strengths. In terms of material lifespan, ferritin and CaCO_3_ each were stable over the measured timeframes (9 & 24 months for ferritin; 15 months for CaCO_3_). Ferritin and CaCO_3_ could therefore be used within susceptibility phantoms to support 3T/7T harmonization studies, using our measurement data as a reference when constructing such a phantom.

**Figure 4:**
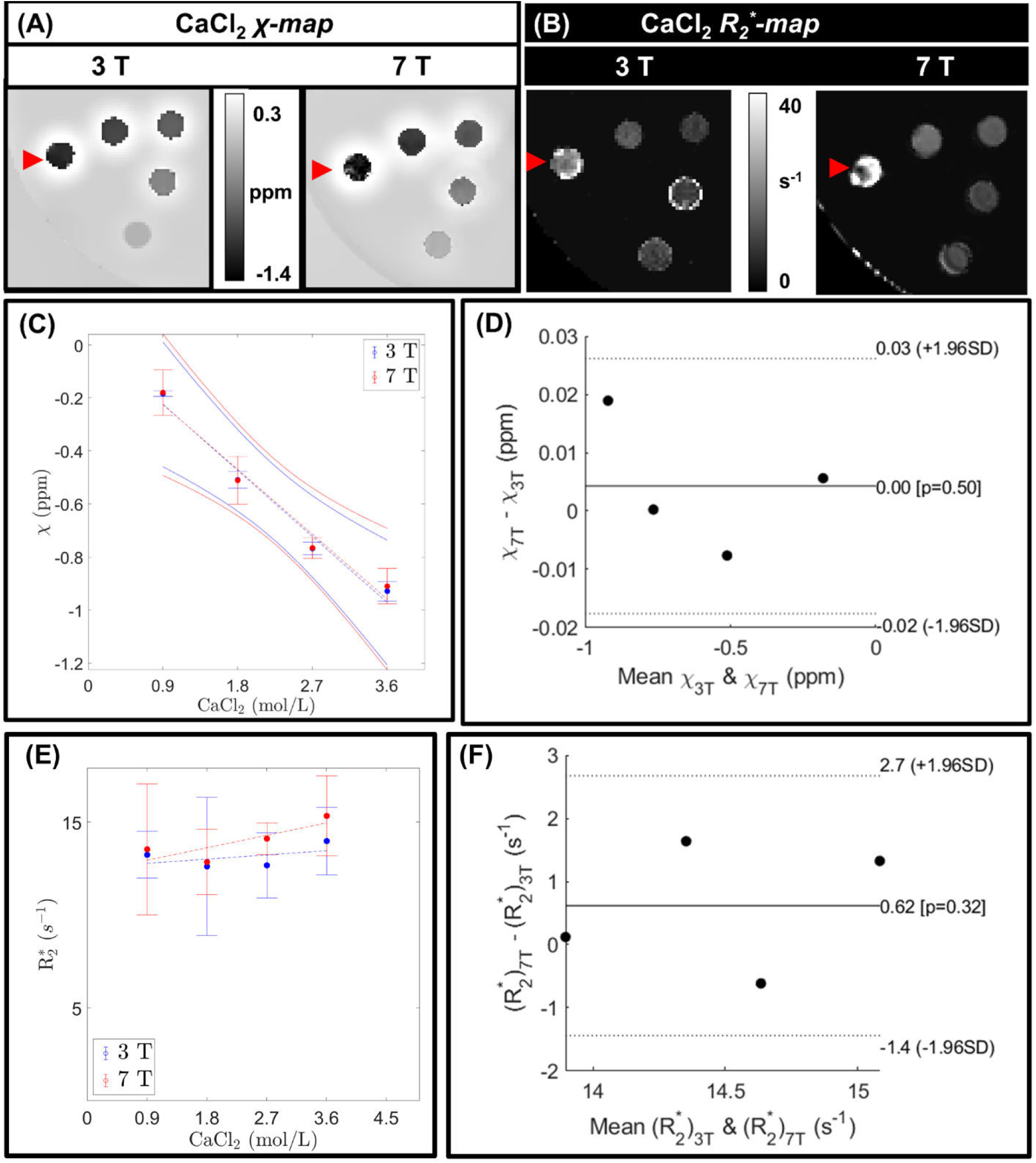
Calcium chloride (CaCl_2_) χ maps (A) and R_2_* maps (B). The voxels corresponding to CaCl_2_ 4.5 mol/L were heterogeneous, as indicated with a red arrowhead on the 7T χ map. (C) Scatter plot of χ as a function of concentration of CaCl_2_. (D) Bland-Altman plot of χ_3T_ versus χ_7T_. (E) Scatter plot of R_2_* as a function of concentration of CaCl_2_. (F) Bland-Altman plot of (R_2_*)_3T_ versus (R_2_*)_7T_.

**Figure 5:**
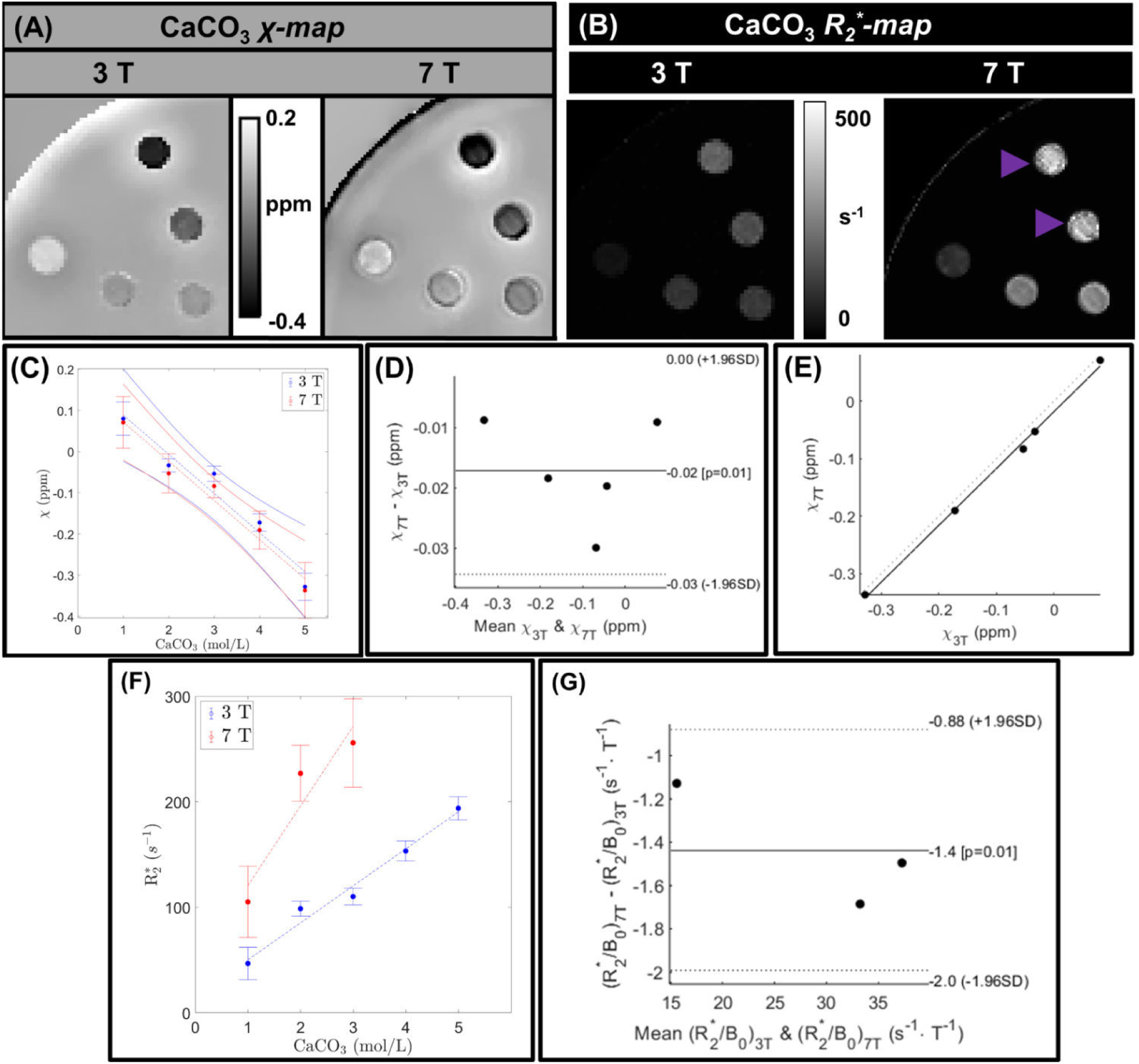
Calcium carbonate (CaCO_3_) χ maps (A) and R_2_* maps (B) of. At 7T, the R_2_* at concentrations ≥ 4.0 mol/L were heterogeneous, indicated with a purple arrowhead. (C) Scatter plot of χ as a function of concentration of CaCO_3_. (D) Bland-Altman and (E) correlation plots of χ_3T_ versus χ_7T_. (F) Scatter plot of R_2_* as a function of concentration of CaCO_3_. (G) Bland-Altman plot of (R_2_*/B_0_)_3T_ versus (R_2_*/B_0_)_7T_.

### 4.3 Cross-field strength agreement

A key motivation of ultra-high-field MRI is the comparability of quantitative measurements of phase and/or susceptibility to clinical MRI^75^. Optimizing and standardizing QSM acquisition and reconstruction protocols across field strength is essential to make susceptibility a robust biomarker. Initiatives such as the German Ultrahigh Field Imaging network traveling heads study^76,77^ and the United Kingdom 7T study^78^ are steps in this direction. Cross-field strength agreement was conducted in the current work using Bland-Altman analysis. For both ferritin and CaCO_3_, the χ_3T_ versus χ_7T_ relationship showed a significant bias; -0.03 ppm (p=0.00) for ferritin and -0.02 ppm (p=0.01) for CaCO_3_. Curiously, the agreement between χ_3T_ and χ_7T_ was best at concentrations 1 and 5 (vials positioned closest to the edge). This could be due to the closer proximity to the receive elements, therefore, resulting in a higher SNR than concentrations 2 to 4 (vials positioned closest to the center). For both ferritin and CaCO_3_, the correlation between χ_3T_ versus χ_7T_ was significant (p=0.02), and the correlation line was y=1.00x-0.03 for ferritin, and y=0.99x-0.02 for CaCO_3_. The consistent cross-field strength bias might provide insight into future 3T/7T harmonization studies. An important property of CaCO_3_ is that the quotient of R_2_* and field strength, R_2_*/B_0_, is invariant with field strength^31^. The correlation between (R_2_*/B_0_)_3T_ and (R_2_*/B_0_)_7T_ was y=0.98x-0.81 (R^2^=1.00). The findings reinforce the use of CaCO_3_ in phantoms for susceptibility source differentiation which require diamagnetic material.

### 4.4 Comparison to reference values

In a USPIO phantom study by Liu et al., it is described that USPIO’s magnetization (χ·B_0_) reaches a saturation point prior to 3T, becoming invariant between 3T and 7T^4^. Liu et al. reported that the USPIO χ_mol_ at 3T; 7T is approximately 1.79; 0.74 ppm·L·mmol^−1 4^. These values fall within the standard errors of χ_mol_ measured in our study, 1.67 ± 0.24; 0.74 ± 0.09 ppm·L·mmol^−1^ at 3T; 7T. Liu et al. reported that the USPIO R_2_*_mol_ at 3T; 7T is 87.2; 106.5 s^-1^·L·mmol^−1 4^. Their data fall within the standard errors of R_2_*_mol_ at 3T (84.2 ± 7.9 s^-1^·L·mmol^−1^), but not at 7T (79.5 ± 4.4 s^-1^·L·mmol^−1^). A possible explanation could be that the USPIO used in this study, Molday Ion, differs from the USPIO used by Liu et al., Ferumoxytol^4^. Previous studies have reported the ferritin susceptibility at 293 K is approximately 0.077 ppm·L·mmol^−1 79^ and 0.080 ppm·L·mmol^−1 80^. These values fall within the range of ferritin χ_mol_ measured in our study, (8.13 ± 1.35)×10^-2^; (8.13 ± 1.19)×10^-2^ ppm·L·mmol^−1^ at 3T; 7T. The susceptibility value of any paramagnetic material may deviate with temperature (by Curie’s Law)^79^, and the internal temperature of the phantom bore frequently deviates from the ambient temperature of the scanner room^72^. Future studies would benefit from using integrated temperature monitoring and/or control^81,82^.

### 4.5 Recommendations

In future studies, we recommend susceptibility matching the undoped agarose to the surrounding fluid by doping each agarose mixture with the same concentration of conductivity modifier used to dope the surrounding fluid. By Equation (6), χ_undoped agarose_= χ_surrounding fluid_ would ensure that χ’_0_ =0. We recommend using ferritin and CaCO_3_ as paramagnetic and diamagnetic susceptibility sources for the validation of QSM imaging

## Conclusion

This research performed a range of quantitative analysis of materials for QSM phantom construction: signal-to-noise ratio, χ and R_2_* maps, outliers, test-retest repeatability, cross-field strength agreement, and material lifespan. Based on the results, we recommend using ferritin and CaCO_3_ as paramagnetic and diamagnetic susceptibility sources for the validation of QSM imaging.

## Acknowledgements

We are grateful to Carl Dixon from the CAI workshop for engineering design input and for machining the phantom. We thank Martijn Cloos for providing a model of the 7T Nova Medical 32 RX-element coil. This work was published in preliminary form^65^. This research was funded by the Australian Government through the Australian Research Council (project number IC170100035). The authors acknowledge the facilities and scientific and technical assistance of National Imaging Facility, a National Collaborative Research Infrastructure Strategy (NCRIS) capability, at the University of Queensland.

## Data availability statement

We facilitate the reproducibility of this study and the phantom by providing scripts (https://github.com/paddyhooper93/Thesis, commit hash 5ea6db1).

