## Supplementary Information for "Evaluation of phantom doping materials in quantitative susceptibility mapping"

### Supporting Information Captions

Table S1: SNR at each TE for each material concentration, at (a) 3T, and (b) 7T, respectively. Low SNR defined as SNR<10 a.u.

Table S2: Clump/agglomerate analysis using $R_2^*$ local outliers at 3 T; 7 T. All values were reported as mean ± standard deviation.

Table S3: Linear fitting coefficients ($c_{mol}$ versus $R_2^*$) and linearity ($R^2$) at 3T; 7T, respectively. Coefficients were reported with standard error.

Figure S1: Reference mask ($M_{reference}$) pipeline: (A) $R_2^*$ thresholding at 5 $s^{-1}$ to separate the vials from the background fluid, indicating $R_2^*$ errors with red arrows. (B) Morphological closing (to fill in small holes within the background fluid, while preserving the larger holes of the vials). (C) Morphological erosion (since cylinders produce an external field affected by the object's susceptibility, we must exclude this external field when referencing). We used a structured spherical element of radius 3 mm and 10 mm for morphological closing and erosion, respectively.

Figure S2: Percentage threshold ($c_\nabla$) versus streaking artifact ($SD_{Ref}$) at (A) 3T, (B) 7T. $SD_{Ref}$ was equal to the standard deviation within the reference mask ($M_{reference}$).

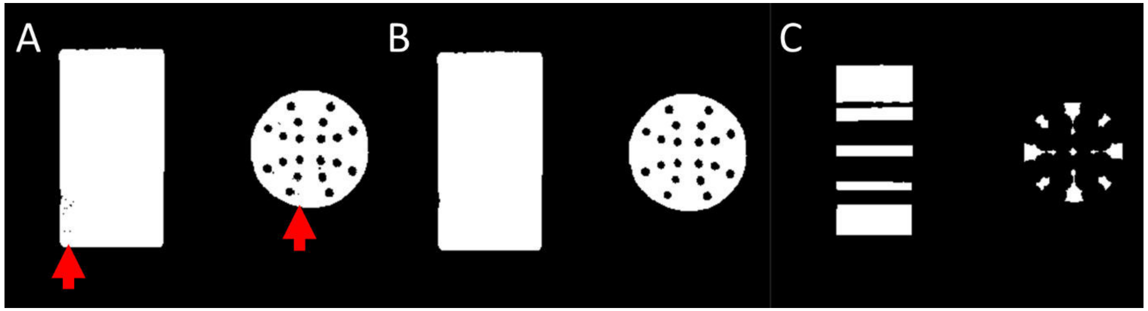


Figure S1: Reference masking ($M_{reference}$) pipeline: (A) $R_2^*$ thresholding at $5\ s^{-1}$ to separate the vials from the background fluid, indicating $R_2^*$ errors with red arrows. (B) Morphological closing (to fill in small holes within the background fluid, while preserving the larger holes of the vials). (C) Morphological erosion (since cylinders produce an external field affected by the object's susceptibility, we excluded this external field when referencing). We used a structured spherical element of radius 3 mm and 10 mm for morphological closing and erosion, respectively.

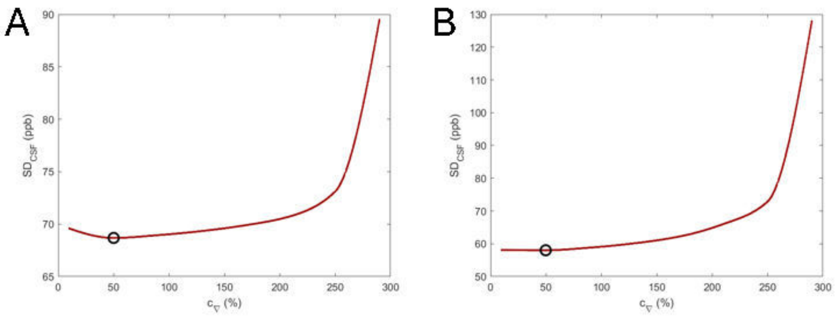


Figure S2: Percentage threshold ($c_\nabla$) versus streaking artifact ($SD_{Ref}$) at (A) 3T, (B) 7T. $SD_{Ref}$ was equal to the standard deviation within $M_{reference}$.

| Table S1 (a) | | TE1: 1.87 | TE2: 3.74 | TE3: 5.61 | TE4: 7.48 | TE5: 9.35 | TE6: 11.22 | TE7: 13.09 | TE8: 14.96 | TE9: 16.83 | TE10: 18.7 | TE11: 20.57 | TE12: 22.44 |
|---|---|---|---|---|---|---|---|---|---|---|---|---|---|
| Fe mmol/L | | | | | | | | | | | | | |
| USPIO | 0.224 | 137 | 125 | 119 | 112 | 106 | 99 | 97 | 91 | 85 | 79 | 77 | 72 |
| | 0.336 | 135 | 122 | 114 | 104 | 97 | 90 | 86 | 78 | 71 | 67 | 62 | 59 |
| | 0.448 | 143 | 127 | 117 | 107 | 96 | 89 | 82 | 75 | 67 | 62 | 56 | 51 |
| | 0.560 | 158 | 137 | 125 | 114 | 103 | 92 | 88 | 78 | 69 | 61 | 57 | 51 |
| | 0.672 | 189 | 161 | 143 | 128 | 112 | 100 | 92 | 80 | 70 | 62 | 56 | 49 |
| Fe mmol/L | | | | | | | | | | | | | |
| Ferritin | 3.76 | 82 | 78 | 76 | 76 | 72 | 70 | 71 | 68 | 65 | 63 | 61 | 60 |
| | 5.37 | 60 | 56 | 55 | 53 | 52 | 51 | 48 | 48 | 45 | 44 | 44 | 43 |
| | 6.98 | 59 | 57 | 53 | 54 | 51 | 49 | 49 | 47 | 45 | 43 | 42 | 41 |
| | 8.60 | 78 | 74 | 70 | 68 | 64 | 63 | 64 | 61 | 56 | 54 | 52 | 52 |
| | 10.21 | 114 | 109 | 102 | 100 | 94 | 93 | 92 | 88 | 82 | 81 | 77 | 76 |
| mol/L | | | | | | | | | | | | | |
| $CaCl_2$ | 0.9 | 67 | 65 | 63 | 64 | 60 | 59 | 60 | 60 | 56 | 55 | 54 | 54 |
| | 1.8 | 46 | 47 | 43 | 45 | 42 | 42 | 41 | 41 | 38 | 38 | 36 | 37 |
| | 2.7 | 49 | 47 | 46 | 47 | 44 | 44 | 44 | 42 | 40 | 39 | 38 | 38 |
| | 3.6 | 66 | 63 | 60 | 59 | 56 | 56 | 54 | 53 | 49 | 50 | 48 | 46 |
| | 4.5 | 91 | 86 | 82 | 82 | 75 | 75 | 72 | 70 | 65 | 63 | 59 | 60 |
| mol/L | | | | | | | | | | | | | |
| $CaCO_3$ | 1.0 | 92 | 89 | 82 | 76 | 72 | 65 | 59 | 54 | 50 | 47 | 43 | 37 |
| | 2.0 | 67 | 59 | 48 | 41 | 34 | 26 | 23 | 19 | 15 | 13 | 12 | 10 |
| | 3.0 | 53 | 46 | 38 | 29 | 25 | 21 | 16 | 14 | 9 | 12 | 6 | 7 |
| | 4.0 | 59 | 46 | 37 | 25 | 20 | 11 | 10 | 9 | 8 | 7 | 5 | 5 |
| | 5.0 | 70 | 54 | 36 | 24 | 18 | 13 | 10 | 8 | 7 | 5 | 6 | 6 |

| Table S1 (b) | | TE1: 3.15 | TE2: 6.3 | TE3: 9.45 | TE4: 12.6 | TE5: 15.75 | TE6: 18.9 | TE7: 22.05 | TE8: 25.2 | TE9: 28.35 |
|---|---|---|---|---|---|---|---|---|---|---|
| Fe mmol/L | | | | | | | | | | |
| USPIO | 0.224 | 154 | 142 | 128 | 116 | 106 | 96 | 87 | 79 | 72 |
| | 0.336 | 160 | 141 | 124 | 108 | 96 | 83 | 74 | 65 | 57 |
| | 0.448 | 173 | 148 | 125 | 105 | 90 | 76 | 66 | 57 | 47 |
| | 0.560 | 159 | 134 | 113 | 93 | 80 | 68 | 57 | 47 | 40 |
| | 0.672 | 139 | 113 | 91 | 73 | 61 | 50 | 40 | 32 | 28 |
| Fe mmol/L | | | | | | | | | | |
| Ferritin | 3.76 | 87 | 82 | 78 | 73 | 69 | 64 | 61 | 58 | 56 |
| | 5.37 | 90 | 84 | 79 | 73 | 70 | 65 | 62 | 58 | 55 |
| | 6.98 | 98 | 90 | 82 | 74 | 70 | 64 | 58 | 54 | 50 |
| | 8.60 | 102 | 92 | 82 | 74 | 68 | 60 | 56 | 50 | 45 |
| | 10.21 | 96 | 85 | 76 | 68 | 62 | 54 | 50 | 44 | 40 |
| mol/L | | | | | | | | | | |
| $CaCl_2$ | 0.9 | 101 | 98 | 95 | 92 | 90 | 87 | 85 | 81 | 79 |
| | 1.8 | 84 | 82 | 78 | 74 | 74 | 72 | 69 | 66 | 64 |
| | 2.7 | 90 | 85 | 79 | 76 | 71 | 68 | 64 | 61 | 57 |
| | 3.6 | 99 | 91 | 87 | 81 | 78 | 74 | 70 | 66 | 63 |
| | 4.5 | 128 | 115 | 100 | 93 | 81 | 77 | 65 | 64 | 53 |
| mol/L | | | | | | | | | | |
| $CaCO_3$ | 1.0 | 58 | 38 | 26 | 18 | 14 | 12 | 10 | 10 | 9 |
| | 2.0 | 43 | 21 | 12 | 9 | 9 | 9 | 9 | 9 | 9 |
| | 3.0 | 38 | 17 | 10 | 9 | 9 | 9 | 9 | 9 | 9 |
| | 4.0 | 28 | 11 | 9 | 9 | 9 | 9 | 9 | 9 | 9 |
| | 5.0 | 21 | 9 | 8 | 9 | 8 | 8 | 8 | 9 | 8 |

Table S1: SNR at each TE for each material concentration, at (a) 3T, and (b) 7T, respectively. The SNR ranges were defined as follows; high SNR (in green): SNR ≥ 50:1, intermediate SNR (in yellow): 20 ≤ SNR < 50, borderline (in orange): 10 ≤ SNR < 20, low (in red): <10 a.u.

| | USPIO | Ferritin | $CaCl_2$ | $CaCO_3$ |
|---|---|---|---|---|
| $R_2^*$ map outlier voxels per vial (%) | 2.65 ±0.65;<br>0.14 ± 0.15 | 1.26 ± 0.35;<br>1.92 ± 0.77 | 1.00 ± 0.07;<br>0.47 ± 0.01 | 0.08 ± 0.09;<br>0.14 ± 0.14 |
| Clump/agglomerate sizes ($mm^3$) | 3.27 ± 3.08;<br>6.40 ± 2.26 | 3.49 ± 3.22;<br>13.01 ± 15.73 | 1.46 ± 0.82;<br>5.59 ± 2.72 | 1.27 ± 0.60;<br>10.40 ± 8.83 |

Table S2: Clump/agglomerate analysis using $R_2^*$ local outliers at 3 T; 7 T. All values were reported as mean ± standard deviation.

| | USPIO | Ferritin | $CaCl_2$ | $CaCO_3$ |
|---|---|---|---|---|
| $R_{2\ mol}^*$ ($s^{-1}\cdot L\cdot mmol^{-1}$) | 84.2 ± 7.9;<br>79.5 ± 4.4 | 0.77 ± 0.16;<br>2.78 ± 0.41 | $10^{-2}\times$ (0.25 ± 0.39);<br>$10^{-2}\times$ (0.74 ± 0.41) | $10^{-1}\times$ (3.52 ± 0.35);<br>$10^{-1}\times$ (8.69 ± 4.99) |
| $R_{2\ 0}^*$ ($s^{-1}$) | 17.0 ± 2.6;<br>16.2 ± 1.1 | 13.3 ± 1.1;<br>8.5 ± 2.6 | 12.5 ± 1.0;<br>12.3 ± 1.0 | 13.8 ± 11.6;<br>45.1 ± 69.1 |
| $R^2$ ($c_{mol}$ versus $R_2^*$) | 0.97; 0.99 | 0.92; 0.93 | 0.22; 0.82 | 0.97; 0.85 |

Table S3: Linear fitting coefficients ($c_{mol}$ versus $R_2^*$) and linearity ($R^2$) at 3T; 7T, respectively. Coefficients were reported with standard error.

| | USPIO | Ferritin | $CaCl_2$ | $CaCO_3$ |
|---|---|---|---|---|
| Bias (ppb) | 8.2 (p=0.24); -3.6 (p=0.46) | 7.9 (p=0.20); -8.0 (p=0.21) | 7.0 (p=0.19); 13.0 (p=0.06) | 4.4 (p=0.06); 1.4 (p=0.38) |
| Correlation line | y=1.00x+0.00 (p=0.02);<br>y=0.97x+0.01 (p=0.02) | y=0.99x+0.01 (p=0.02);<br>y=0.97x+0.01 (p=0.02) | y=0.99x+0.00 (p=0.02);<br>y=0.97x-0.02 (p=0.02) | y=0.99x+0.00 (p=0.02);<br>y=0.98x+0.00 |

Table S4: Bland-Altman and correlation analysis of $\chi_{analytical}$ versus $\chi_{MEDI+0}$ at 3T; 7T, respectively. The results of a t-test were shown for bias and correlation (null hypotheses: zero bias, no correlation).
